# Oct4 clusters promote DNA accessibility by enhancing chromatin plasticity

**DOI:** 10.1101/2025.10.20.683403

**Authors:** Jan Huertas, M. Julia Maristany, Rosana Collepardo-Guevara

## Abstract

Pioneer transcription factors are defined by their ability to engage closed chromatin and render it accessible. Oct4, a master regulator of pluripotency, exemplifies this capacity as it can bind nucleosome-occupied DNA. Yet, how Oct4 reshapes chromatin fibres beyond the scale of a single nucleosome has remained unclear. Here, we harness our near-atomistic coarse-grained chromatin model to dissect how full-length Oct4 interacts with chromatin fibres of varying nucleosomal repeat lengths. We find that Oct4 increases DNA accessibility not by decondensing chromatin, but by driving fibres into compact liquid-like states in which nucleosomes breathe, reorient, exchange neighbours, and transiently expose DNA. Oct4 binds preferentially to linker DNA through its structured POU domains, while its disordered activation domains mediate cluster formation: a three-stage process consisting of nucleation of small clusters, attachment to chromatin via the DNA binding domains, and growth that is amplified when long DNA linkers enable fibre bending and looping. Oct4 clusters deform chromatin and bridge distal DNA segments in a stoichiometry- and linker length–dependent manner, revealing a tunable mechanism for controlling higher-order structure. Together, our results establish how Oct4 exploits multivalency, nucleosome breathing, and chromatin flexibility to reorganise fibres into liquid-like states that reconcile condensation with enhanced accessibility, providing a physical route to overcome the transcription-factor ‘search problem’ and activate silenced genes during reprogramming.

## Introduction

In eukaryotic cells, nearly two metres of DNA encoding the genetic information of the cell are confined within a nucleus of ≈10*µ*m in diameter. This remarkable level of compaction is achieved through the assembly of DNA into chromatin, a nucleoprotein polymer whose spatiotemporal organisation is not random but tightly regulated^[1,2]^. Chromatin structure is increasingly recognised as a dynamic, context-dependent regulator of DNA accessibility that both modulates transcription factor binding to control gene expression and adapts in response to transcriptional activity^[3–5]^.

The basic unit of chromatin is the nucleosome, a complex of approximately 147 base pairs of DNA wrapped around a histone octamer^[6]^. Nucleosomes are joined by linker DNA segments of varying lengths, resulting in a 10-nm ‘beads-on-a-string’ fibre^[7]^. Each nucleosome contains two copies each of histones H2A, H2B, H3, and H4, together with ten intrinsically disordered histone tails (the N-termini of all four histones and the short C-termini of the two H2A proteins)^[8]^. These highly charged, flexible tails serve as hotspots for post-translational modifications^[9,10]^. Histone tails mediate interactions with DNA, other nucleosomes, and chromatin-associated proteins, providing an enthalpic gain that counteracts the electrostatic repulsion of DNA and promoting compaction^[11–15]^.

Early models proposed that the 10-nm fibre folds into a rigid 30-nm zigzag fibre, stabilised by regular second-nucleosome neighbour interactions^[16–19]^. However, advances in live-cell imaging and cryo-electron tomography^[20–23]^ have instead revealed that chromatin adopts an irregular, dynamic, and heterogeneous organisation—also termed ‘liquid-like’ to emphasize the lack of long-range translational order^[22,24,25]^. The dynamic liquid-like behaviour of chromatin is sustained by short-lived multivalent nucleosome–nucleosome interactions that are underpinned by weak, promiscuous, and transient DNA–histone tail contacts^[15,26]^. Consistently, nucleosomes in cells reconfigure within heterogeneous clusters of variable density^[27,28]^, functional states are distinguished more by nucleosome packing than by rigid 30-nm order^[19,27,29]^, and chromatin arrays undergo liquid–liquid phase separation ^[15,30–32]^.

Understanding how pioneer transcription factors operate within such a dynamic, irregular and heterogeneous chromatin structural scenario is therefore critical. Chromatin compaction and phase separation jointly contribute to the formation of closed chromatin, in which tight nucleosome–nucleosome contacts restrict access of the transcriptional machinery to DNA. Yet, despite this barrier, pioneer factors can recognise and bind their target DNA sites within nucleosome-occupied regions^[33–35]^. Although the mechanistic basis of this ability remains incompletely understood, pioneer-factor binding to closed chromatin is a key step in gene regulation, particularly during cell-fate transitions^[36,37]^.

Among these, Oct4 is one of the best-characterised pioneer transcription factors and serves as a paradigm for how multidomain proteins engage and remodel compact chromatin^[38–41]^. Oct4 is a 360-amino-acid protein containing a bipartite POU DNA-binding domain flanked by intrinsically disordered activation domains^[42,43]^. It acts as a master regulator of pluripotency: together with Sox2 and Nanog, it forms the core transcriptional circuitry that maintains stem-cell identity^[44]^. As one of the four Yamanaka factors (Oct4, Sox2, Klf4, and c-Myc)^[45]^, Oct4 is sufficient to reprogramme somatic cells into induced pluripotent stem cells (iPSCs). Notably, three of these factors (Oct4, Sox2, and Klf4) are pioneer transcription factors^[46,47]^, underscoring that the ability to engage closed chromatin is crucial for cellular reprogramming^[48]^.

The intrinsically disordered and structured domains of Oct4 enable multivalent self-interactions and interactions with other nuclear components such as Mediator subunits and DNA, driving its incorporation into transcriptional condensates^[49–51]^. These Oct4-containing condensates are thought to concentrate transcriptional machinery, promote enhancer–promoter proximity, and thereby modulate gene expression programs^[52]^. More broadly, biomolecular phase separation is now recognised as a fundamental organising principle of the nucleus, influencing both the compaction of DNA into heterochromatin domains^[53]^ and the local enrichment of transcriptional regulators at active genomic loci^[50]^. Within this framework, condensates selectively concentrate subsets of proteins and chromatin, while providing physicochemical environments (defined by their thermodynamic parameters and material properties) that can regulate chemical reactions and transcriptional output. Consistent with this view, transcriptional condensates have been shown to modulate inflammatory gene expression^[51]^, and HP1*α* condensates drive gene silencing^[54]^.

Importantly, the relationship between protein condensates and chromatin is reciprocal: chromatin provides an extended scaffold with multiple DNA binding sites that can lower nucleation thresholds and localise condensates to specific genomic loci^[55,56]^. In addition to phase separation, protein–DNA interactions can also drive surface condensation through pre-wetting transitions, in which proteins condense on DNA below their homotypic saturation concentration^[55,57]^. A striking example is Klf4, which undergoes sequence-dependent condensation on DNA through such a mechanism^[57]^.

Uncovering the mechanistic principles by which transcription factors recognise and engage chromatin with submolecular resolution remains a major challenge. Cryo-EM has resolved only a limited set of pioneer factor–nucleosome complexes^[58]^, which predominantly correspond to enthalpically stabilised conformations and systematically underrepresent the intrinsically disordered domains that confer structural plasticity. For Oct4, several nucleosome-bound structures have been reported^[39–41]^, yet these correspond to isolated low-energy minima on a rugged free-energy landscape. Furthermore, the available structures often resolve only one half of the bipartite POU domain. Such static minima provide atomic detail but cannot capture the dynamic fluctuations, transiently unwrapped states, contributions of disordered domains, and transition pathways that underpin the function of pioneer factors. Molecular Dynamics (MD) simulations can complement experiments by capturing the full dynamical ensembles sampled by the protein–nucleosome complexes, the probabilities of each configuration, and the free-energy barriers connecting them. For single nucleosomes, our atomistic simulations quantified the stability of Oct4 to nucleosome binding, identified the residues mediating key protein–DNA contacts^[59]^, and revealed how the bipartite POU domain biases the free-energy landscape towards partially unwrapped nucleosomal states^[38]^.

Atomistic MD simulations of chromatin assemblies remain computationally intractable, motivating the use of coarse-grained approaches^[60,61]^. Here, we employ a chemical-specific multiscale chromatin model^[26]^ that explicitly represents amino acids and DNA base pairs, thereby retaining residue-level physicochemical properties while enabling chromatin simulations. This framework has successfully reproduced cryo-ET reconstructions of chromatin condensates at near-atomistic resolution^[15]^, probed nucleosome dynamics at genomic loci^[62]^, and captured the experimentally-observed liquid-like behaviour of linker histone H1 within chromatin domains in human cells^[63]^.

Pioneer transcription factors such as Oct4 are classically thought to ‘open’ chromatin, yet how they remodel chromatin beyond a single nucleosome remains poorly understood. Using chemical-specific coarsegrained molecular dynamics simulations of full-length Oct4 bound to 12-nucleosome fibres, we show that Oct4 enhances chromatin accessibility not by global decondensation but by promoting nucleosome breathing within a compact, liquid-like fibre. Oct4 multivalency drives clustering and fibre reorganisation through a three-stage mechanism involving nucleosome breathing, chromatin bending, and cluster coalescence. Together, these findings reveal how Oct4 leverages multivalency, nucleosome breathing, and chromatin flexibility to generate DNA accessibility within compact fibres, providing a route to alleviate the transcription-factor ‘search problem’. This establishes a physical framework for pioneer-factor action and provides testable predictions linking chromatin architecture, Oct4 dosage, and reprogramming efficiency..

## Results

### Concentration-dependent liquid-like reorganisation of chromatin by Oct4

To dissect how Oct4 influences chromatin structure, we simulated 12-nucleosome fibres with a nucleosomal repeat length (NRL = 147 bp of nucleosomal DNA + linker DNA length) of 200 bp, which is within the values frequently found in heterochromatic regions of higher-eukaryotes^[64,65]^. We used our chemically specific chromatin model^[26]^, in which amino acids, DNA base pairs, and phosphate charges are explicitly represented (Fig. 1A and B). Full-length Oct4 was coarse-grained at one bead per residue, maintaining the structure of the globular DNA-binding domains (POU_S_ and POU_HD_) via rigid bodies while treating all intrinsically disordered regions as flexible polymers (see Methods). This resolution retains physicochemical detail while enabling efficient sampling of Oct4 and chromatin.

**Fig. 1.**
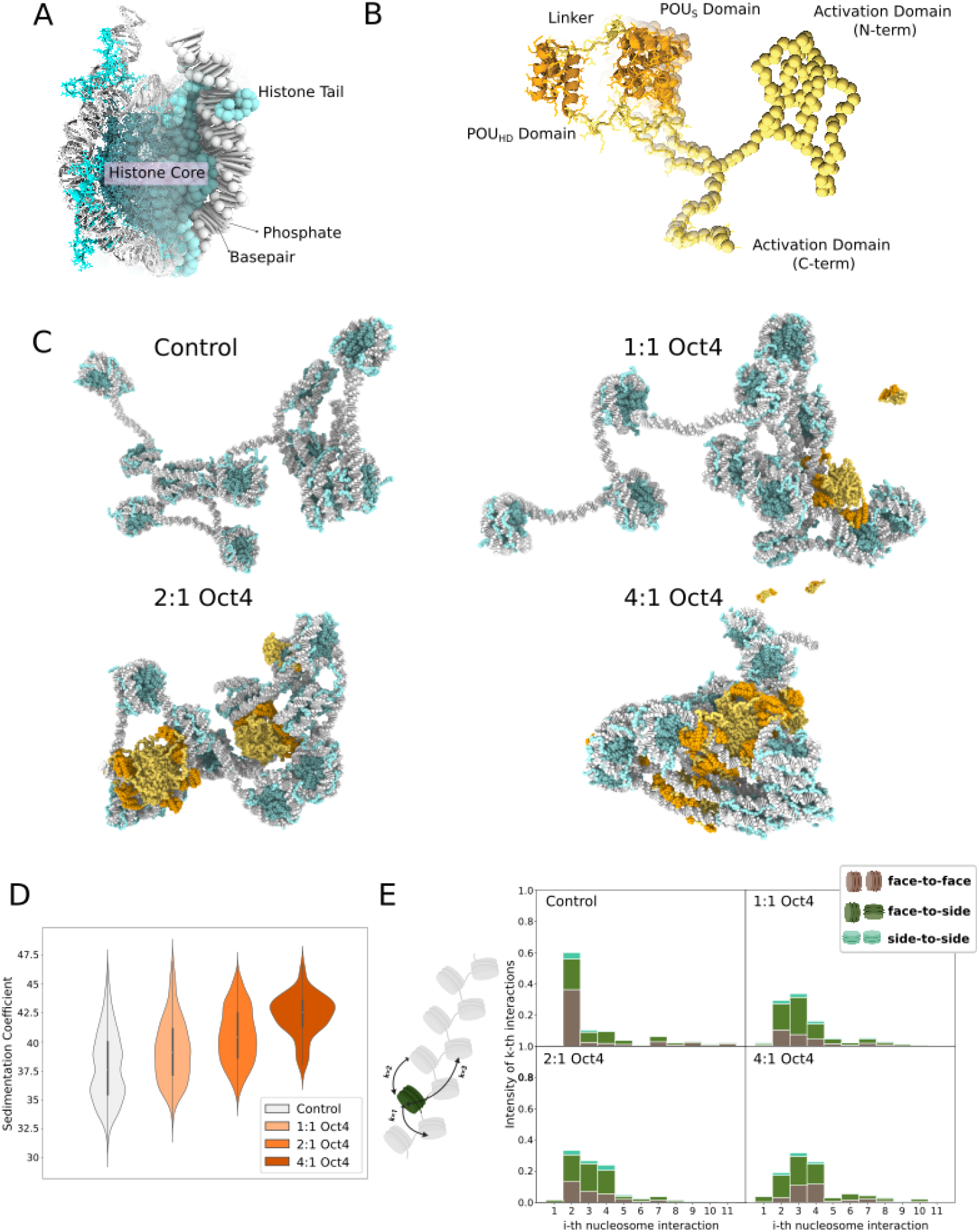
Coarse-grained simulations of chromatin reveal an Oct4-dependent chromatin compaction. **A**. Our chromatin model, where every amino-acid is represented as a single bead and every basepair of DNA as an ellipsoid with two charged phosphates. The secondary structure of the histone core is held via an elastic network model. **B**. Model of Oct4, coarse-grained at the same resolution as histones. The secondary structure of the DNA-binding domains is also kept via the use of a Gaussian elastic network model. **C**. Representative snapshots of a chromatin fibre of 12 nucleosomes with increasing Oct4:Nucleosome stoichiometries. **D**. Sedimentation coefficient of chromatin with different Oct4:Nucleosome stoichiometries. **E**. Frequency of interactions among k-th nearest nucleosome neighbours for chromatin with different Oct4:Nucleosome stoichiometries. The bars are colour-coded to represent the percentage of nucleosome pairs involved in different types of interactions: face-to-face (brown), face-to-side (dark green), and side-to-side (turquoise).

Our simulations revealed that Oct4 compacts chromatin in a concentration-dependent manner (Fig. 1C), as reflected by a monotonic increase in the sedimentation coefficient (Fig. 1D), from a median of 37.6 S (in simulations with 0:1 Oct4:nucleosome ratio) to 42.5 S (4:1 Oct4:nucleosome ratio). At low stoichiometry (1:1), isolated Oct4 molecules perturb nucleosomes locally but cannot reorganise the fibre globally. As Oct4 concentration increases, multivalent weak and transient Oct4–Oct4 and Oct4–DNA interactions nucleate clusters on chromatin, which act as hubs bridging distal nucleosomes and generating a network of transient contacts. Akin to condensate growth, Oct4 clusters expand as additional molecules are recruited into the multivalent interaction network. In this system, multivalency arises from the combination of the Oct4 disordered activation domains, the bipartite DNA-binding domains, and the abundance of chromatin binding sites.

At first glance, chromatin compaction driven by Oct4 may seem counterintuitive, as pioneer transcription factors are typically defined by their ability to engage closed chromatin and make it accessible to other factors, which is often equated with fibre decompaction. Indeed, pioneer transcription factors are among the few proteins capable of directly binding nucleosome-occupied DNA within compact chromatin. Yet, our simulations uncover a striking cooperative effect: chromatin compaction and DNA accessibility are not contradictory, but can be jointly favoured by the same underlying thermodynamics. We found that Oct4 enhances DNA accessibility while condensing chromatin because it promotes nucleosome breathing and drives chromatin into a liquid-like state^[26,53]^. By liquid-like, we refer to a configurational ensemble lacking long-range translational or orientational order, in which nucleosomes continuously reorient, exchange neighbours, and explore a broad spectrum of disordered configurations^[26]^. This liquid-like state is reflected in the reorganisation of inter-nucleosome contact topology (Fig. 1E): in Oct4-free fibres, internucleosomal contacts are dominated by long-lived second-neighbour face-to-face interactions (i.e., zig-zag contacts), whereas Oct4 binding promotes transient, heterogeneous contacts across multiple neighbours. Live-cell single-molecule tracking experiments confirm this behaviour in vivo: Oct4 binds preferentially within flexible, viscoelastic chromatin microenvironments^[66]^.

Mechanistically, Oct4 molecules enable a liquid-like chromatin state by promoting: (i) Nucleosome breathing: Oct4 binding to the nucleosome, targets the DNA at the entry/exit site, allowing breakage of histone–DNA contacts in that region and, thus, stabilising the partially unwrapped nucleosome intermediates, reducing the unwrapping barrier and enriching the ensemble with partially unwrapped states^[38,40,41,59,67,68]^. This occurs because the Oct4–DNA binding partially compensates for the enthalpic penalty associated with the loss of histone–DNA contacts during unwrapping. At the same time, the increased DNA fluctuations during breathing expand the accessible conformational phase space of the nucleosome, conferring an entropic gain that further stabilises transiently unwrapped states. (ii) Linker DNA bending and torsional fluctuations: Oct4 binding lowers the energetic penalty for DNA bending and torsion, facilitating linker DNAs to adopt a broader range of conformations. This is evident in the increased fraction of first-neighbour interactions observed in our simulations (Fig. 1E). (iii) Inter-nucleosome neighbour exchange: By binding in between nucleosomes and promoting breathing, Oct4 disrupts the ordered face-to-face zigzag contacts and stabilises heterogeneous ones, thereby lowering the barrier to neighbour exchange. By simultaneously reducing these barriers, Oct4 enables chromatin to explore a broad ensemble of disordered states.

From a thermodynamic point of view, enabling DNA accessibility by forming a compact liquid-like chromatin state is preferred over decompaction. To decompact chromatin, Oct4 binding would have to offset the substantial enthalpic cost of disrupting nucleosome–nucleosome interactions. In contrast, driving chromatin into a liquid-like state preserves the enthalpically favourable nucleosome–nucleosome contacts while simultaneously increasing the configurational entropy of chromatin, thus lowering the free energy relative to both a rigid zigzag and a decompacted fibre. Therefore, Oct4 renders chromatin accessible by reorganising it into a compact, liquid-like state, as that represents the thermodynamically optimal solution.

### Linker DNA length tunes Oct4-mediated chromatin remodelling

Having established that Oct4 can drive chromatin into a compact liquid-like state, we next asked whether structural variations of the chromatin fibre modulate its effect. A key parameter regulating chromatin structure is the nucleosomal repeat length, which determines both the spacing and relative orientations of nucleosomes along the fibre. Because the DNA helix twists by 360^*°*^ every ∼10.5 bp^[69]^, a 5 bp difference in linker length rotates the entry–exit DNA by nearly half a helical turn, thereby drastically reorienting adjacent nucleosomes^[15,31]^. With 10*n* linkers, nucleosomes align in parallel orientations, favouring regular zig-zag structures stabilised by face-to-face nucleosome stacking^[15,31]^. By contrast, with 10*n*+5 linkers, nucleosomes adopt anti-parallel orientations, which frustrates face-to-face stacking and promotes structural disorder^[15,31]^.

To examine how linker DNA length modulates the ability of Oct4 to reorganise chromatin, we simulated 12-nucleosome fibres with linker lengths of 20 bp (NRL = 167 bp), 25 bp (172 bp), 30 bp (177 bp), and 35 bp (182 bp), thereby enabling a direct comparison between 10*n* and 10*n*+5 periodicities. In these simulations, Oct4 compacted all fibres with long DNA linkers (≥30 bp; NRLs of 177, 182, and 200 bp), had little impact at intermediate length (25 bp; NRL = 172 bp), and decompacted fibres with short linkers (20 bp; NRL = 167 bp; Fig. 2A). For long DNA linkers, DNA bending deformations can be distributed over an extended DNA contour, lowering their energetic cost relative to shorter linkers. This facilitates the formation of DNA loops and long-range contacts, while the additional DNA length also presents more potential binding sites for Oct4. Together, these effects enable Oct4 to bridge distal segments and promote fibre compaction (Fig. 2B, Supplementary Fig. S1A). At the intermediate linker length of 25 bp (NRL = 172 bp), the linkers are too short to support bridging interactions, explaining the lack of Oct4-driven compaction. In contrast, in the absence of Oct4, the short linker length of 20 bp (NRL = 167 bp) favours the canonical zigzag configuration, with nucleosomes stacked face–face and straight linkers layered in parallel. This zigzag arrangement is already maximally compact, as the short linkers severely restrict the conformational space accessible to the fibre. Consequently, in the short-linker zigzag, Oct4 must intercalate to reach its DNA-binding sites, disrupting the dense packing and thereby swelling the fibre (Fig. 2B, left).

**Fig. 2.**
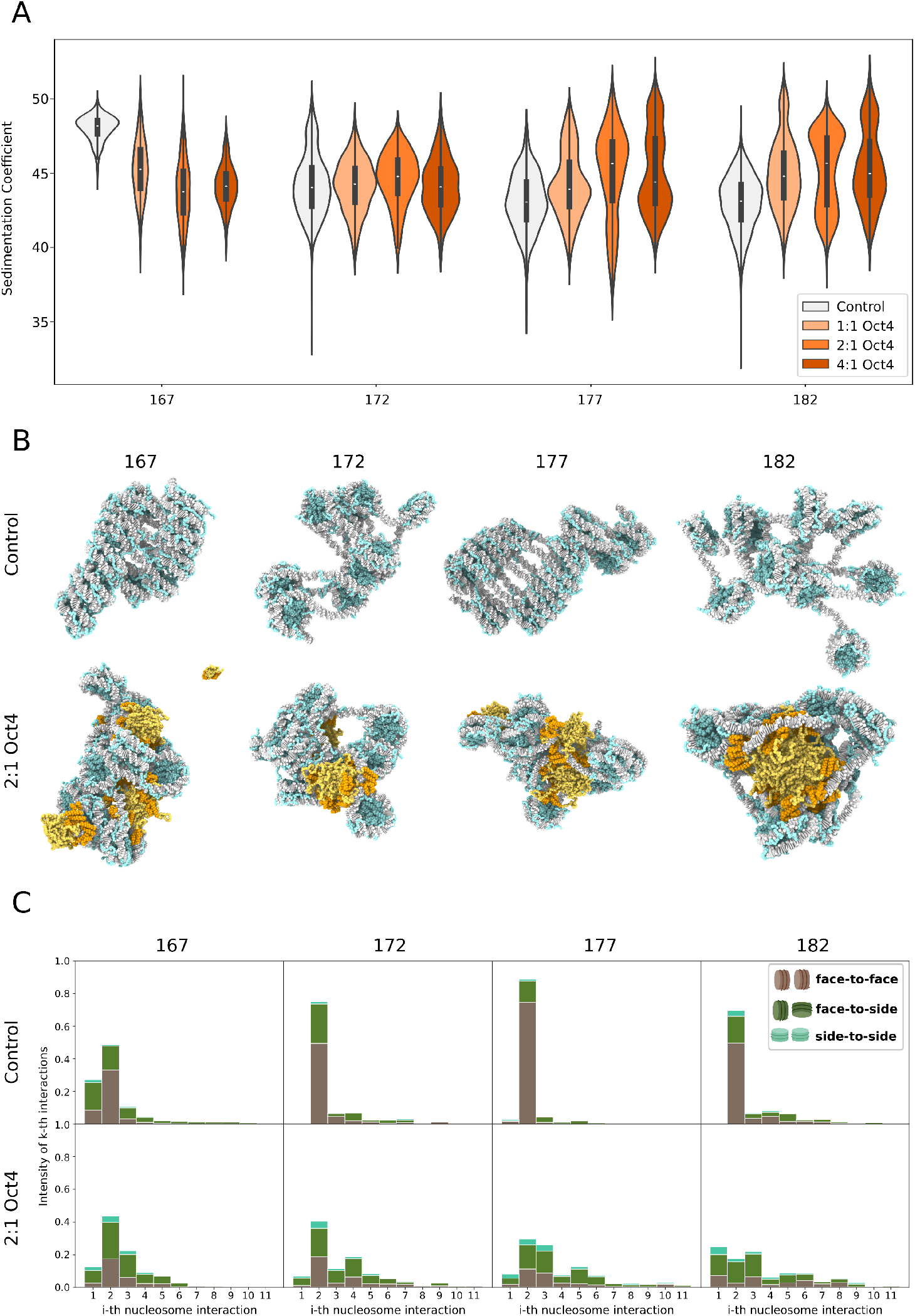
Nucleosomal Repeat Length modulates chromatin’s response to Oct4 binding. **A**. Sedimentation coefficients of chromatin across different NRLs and Oct4:nucleosome stoichiometries. **B**. Representative snapshots of chromatin with varying NRL, in the absence of Oct4 and at a 2:1 Oct4:nucleosome stoichiometry. **C**. Frequency of interactions among k-th nearest nucleosome neighbours for chromatin with different NRLs, in the absence of Oct4 and at a 2:1 Oct4:nucleosome stoichiometry. The bars are colour-coded to represent the percentage of nucleosome pairs involved in different types of interactions: face-to-face (brown), face-to-side (dark green), and side-to-side (turquoise).

Regardless of linker length, Oct4 drove chromatin towards liquid-like behaviour, as evidenced by a marked reduction in stable second-neighbour zigzag interactions and a corresponding increase in heterogeneous inter-nucleosome contacts (Fig. 2C, Supplementary Fig. S1B). This unifying behaviour explains the striking loss of 10*n* versus 10*n*+5 periodicity described above: Oct4 enhances nucleosome breathing, leading to dynamic fluctuations of linker length that effectively interconvert between 10*n* and 10*n*+5 values. As linkers transiently lengthen or shorten, their helical twist also shifts, continuously altering the preferred orientations they impose on neighbouring nucleosomes. As a result, NRL-imposed orientations are replaced by rotational diversity, such that both 10*n* (initially ordered) and 10*n*+5 (already disordered) fibres converge towards Oct4-driven, liquid-like ensembles. In this Oct4-driven liquid-like regime, chromatin explores a rugged free-energy landscape with multiple shallow minima, overriding fibre parameters that would otherwise stabilise ordered states and amplifying disorder where it is intrinsic.

Overall, our findings show that Oct4-mediated chromatin reorganisation is governed by linker DNA length: long linkers enable compaction through bridging, while short linkers force intercalation and decompaction. In both cases, the outcome is a liquid-like state in which fibre condensation and DNA accessibility coexist, underscoring how chromatin architecture modulates the regulatory capacity of pioneer transcription factors.

### Oct4 preferentially binds free linker DNA through its DNA-binding domains

To uncover the mechanistic basis of how Oct4 remodels chromatin, we asked which domains of the protein itself mediate its interactions with linker and nucleosomal DNA in chromatin. To this end, taking advantage of the near-atomistic resolution of our model, we quantified the frequencies of Oct4–DNA contacts across the fibre, breaking them down by Oct4 domains.

We found that while the majority of Oct4–DNA contacts occurred within linker regions (20 to 60% of molecules bound), Oct4 also contacted the nucleosomal DNA (median ≈5-10% of molecules across all NRLs). A small but significant number of Oct4 molecules (2 to 5%) were found to bind to the nucleosome entry-exit (Fig. 3A). The preference of Oct4 to target linker versus nucleosomal DNA was regulated by linker DNA length: fibres with the shortest linkers (20 bp) presented a significantly higher fraction of molecules bound to the nucleosomal DNA (median ≈9%, compared to ≈5–6% for longer linkers). This shift likely reflects both the reduced abundance of free linker DNA in the 20 bp system, and its decreased accessibility within the compact zigzag geometry, which together increase the probability that Oct4 engages nucleosomal sites.

**Fig. 3.**
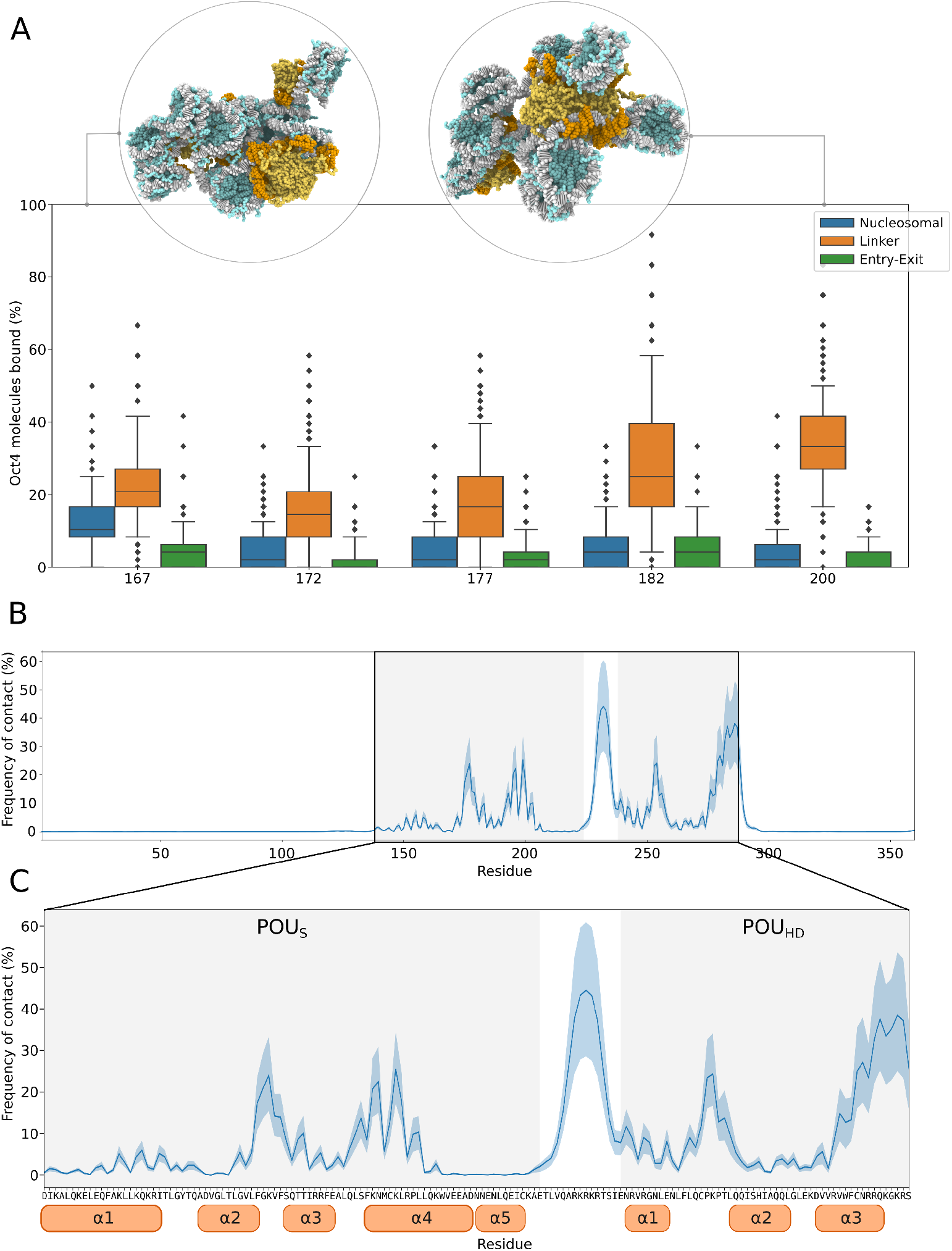
Oct4 binds preferentially to linker DNA via its DNA binding domain. **A**. Percentage of Oct4 molecules bound to the nucleosomal, linker DNA, or the nucleosome entry-exit site, across different NRLs. In inserts, snapshots highlighting interactions with nucleosomal DNA (in NRL 167) or with linker DNA (in NRL 200). **B**. Per residue contact map of the Oct4-DNA interactions, averaged across all NRLs in simulations with 2:1 Oct4 nucleosome stoichiometry. For each residue, the percentage of interaction with a DNA base pair is depicted. The shadowed area represents the standard deviation. **C**. Zoom-In of the protein-DNA contacts for the DNA binding domain shown in panel B, with the amino acids represented in the x-axis. In orange, the helices of the POU_S_ and POU_HD_ domains.

Leveraging the near-atomistic resolution of our model, we explore the binding modes of Oct4 to DNA. We found that DNA binding was overwhelmingly mediated by Oct4’s bipartite DNA-binding domain (DBD), with residues outside the DBD rarely making contacts (Fig. 3B, Supplementary Fig. S2). In our model, Oct4 interacts with DNA without explicit base-specific recognition; any DNA sequence dependence arises from the distinct mechanical deformations associated with different DNA sequences, rather than through direct protein–nucleotide hydrogen bonding. Nonetheless, our simulations recapitulate the experimentally observed binding interfaces of Oct4 to DNA; the helices *α*_4_ of the POU_S_ domain and *α*_3_ of the POU_HD_ domain carried the majority of contacts (peaks of ≈30% and ≈55%, respectively), aligning with observations from crystal structures of Oct4 bound to its specific DNA recognition site^[43]^ (Fig. 3C). This agreement likely emerges from the intrinsic shape and electrostatic complementarity of the POU_S_ and POU_HD_ domains, together with the balanced amino acid–DNA interaction parameters in our model. Interestingly, the flexible linker region of the DBD, modelled here as a disordered region, also formed substantial contacts with DNA (median frequency of contact of ≈45% of the simulation) (Fig. 3C). This observation provides a mechanistic explanation for previous findings that the Oct4 linker region is critical for cellular reprogramming^[43]^, suggesting that its DNA contacts actively modulate Oct4’s pioneer function. More broadly, it illustrates how pioneer transcription factors can exploit both structured and disordered regions to engage chromatin in ways that are tuned by fibre architecture.

### Chromatin acts as a nucleation site for Oct4 clusters

Initial qualitative analysis of our chromatin–Oct4 simulations revealed the formation of Oct4 clusters across all conditions (Fig. 4A). Given that Oct4 has been shown to bind cooperatively to chromatin—often in a hierarchical, context-dependent fashion at enhancers^[70]^—we reasoned that the formation of Oct4 clusters in our simulations potentially reflects a functionally relevant mode of chromatin organisation. To determine the mechanism of Oct4 cluster formation, we performed additional control simulations of Oct4 molecules in the absence of chromatin, with each simulation probing the behaviour of either 12, 24, or 48 Oct4 molecules.

**Fig. 4.**
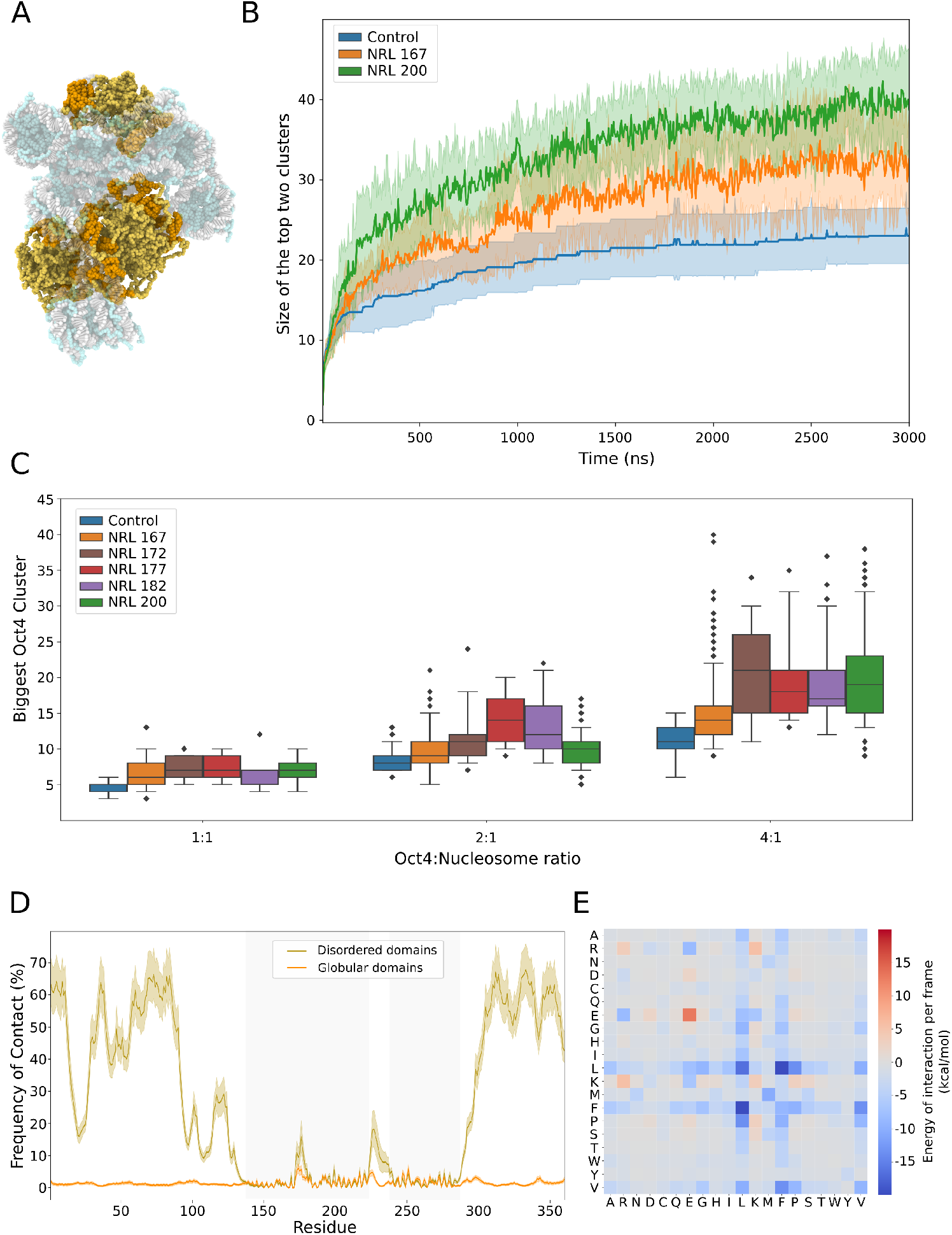
Oct4 forms clusters via IDR contacts that grow in the presence of chromatin. **A**. Snapshot highlighting the formation of Oct4 clusters upon binding to chromatin. **B**. Mean and standard deviation of the sum of the size, in number of molecules, of the two most populated clusters across 10 independent replicas, for the case without chromatin (blue), and NRLs of 167 (orange) and 200 (green). **C**. Size in number of molecules of the biggest Oct4 cluster, in simulations of Oct4 alone (blue) and with varying NRL (167, orange; 172 brown; 177, red; 182, purple; 200, green), across Oct4:Nucleosome ratios. **D**. Per residue contact map of the Oct4-Oct4 interactions, averaged across all NRLs in simulations of 2:1 Oct4 nucleosome stoichiometry. For each residue, the percentage of interaction with a residue belonging to a disordered domain or to a globular domain is depicted. The shadowed area represents the standard deviation. Contacts with disordered domains are represented in gold, and with the globular domains, in orange. **E**. Normalised intermolecular amino acid–amino acid interaction energy matrix. Amino-acid contacts are averaged over all frames and weighted by pair interaction energy and residue abundance to correct for compositional bias.

To quantify the influence of chromatin on Oct4 self-association, we define the cluster size *Q* as the number of Oct4 molecules within a prescribed cutoff distance of one another at a given simulation frame. We monitored the temporal evolution of the two largest clusters, *Q*_1_ + *Q*_2_, as this combined measure is less sensitive to transient fusion and fission events that characterise the early stages of condensation^[55]^. Analysis of these trajectories (Fig. 4B, Supplementary Fig. S3) reveals that chromatin does not significantly accelerate cluster nucleation, as the formation of the first small clusters is observed at similar timescales in all systems. Instead, we observe that chromatin stabilises and promotes the coalescence of existing clusters. While the nucleation of Oct4 clusters appears insensitive to the characteristics of chromatin, their subsequent growth depends strongly on the amount of linker DNA (but not on the 10*n* vs 10*n*+5 periodicity), mirroring the compaction behaviour. By reducing the entropic cost of encounters between small Oct4 assemblies and acting as a bendable scaffold that favours their merging, the chromatin fibre facilitates the growth of clusters that would otherwise remain small.

Oct4 formed clusters even in the absence of chromatin; however, these remain markedly smaller than those observed when chromatin is present (Fig. 4C, Supplementary Fig. S4). In the control simulations (without chromatin), the median size of the largest Oct4 cluster (*Q*_1_) was approximately 5 molecules at a total of 12 Oct4 proteins, 9 molecules at 24, and 12 molecules at 48. In the presence of chromatin, where these correspond to 1:1, 2:1, and 4:1 Oct4:nucleosome stoichiometries, median cluster sizes increased to roughly 6–9 molecules at 1:1, 12–16 at 2:1, and 18–25 at 4:1 (across nucleosome repeat lengths), with upper values exceeding 35–40 molecules per cluster. The differences were evident even at low stoichiometry (12 Oct4 molecules), supporting the view that chromatin promotes the stability of Oct4 clusters.

Cluster size, at equilibrium, scaled with Oct4 concentration, as expected, and was strongly influenced by the amount of linker length, but not by 10*n* vs 10*n*+5 periodicity. Instead, cluster growth followed the same linker length range-dependent trends observed for compaction: short, medium, and long linkers produced distinct outcomes. At short linkers (167 bp), where the fibre is highly compact and rigid, Oct4 clusters could only wet the fibre surface, resulting in modest growth (≈6 to 11 molecules from 1:1 to 4:1). At intermediate linkers (172 bp), the additional DNA length provided more accessible binding sites, enabling moderate cluster growth. At long linkers (177–200 bp), Oct4 not only had more DNA to engage but could also bend and loop the fibre (as discussed in the previous section), allowing chromatin to wrap around and bring together small clusters to promote their coalescence; this interplay of increased DNA binding sites and enhanced fibre flexibility produced the largest clusters (≈7 to 19 molecules).

To characterise how those clusters were formed, we computed the protein–protein interactions of Oct4 at the amino acid level (Fig. 4D, Supplementary Fig. S5). Remarkably, most contacts were mediated by the intrinsically disordered activation domains of Oct4. Residues within disordered regions of Oct4 are involved in mediating Oct4–Oct4 contacts for a median of around 60% of the trajectory, whereas residues in the globular domains were typically involved 3%, with only some residues around 15%. This behaviour was also observed in the simulations without chromatin (Supplementary Fig. S5A to C). Further analysis revealed that cluster stability was significantly contributed by hydrophobic associations among leucine (L) and phenylalanine (F), with L–L, L–F and F–F pairs showing the strongest attractions (mean interaction energy per frame with values of -10 to -15 kcal/mol; Fig. 4E). In contrast, the high content of glutamic acid in the N-terminal activation domain (14 out of 137 residues) results in electrostatic repulsion that frustrated the growth of the Oct4 clusters in the absence of chromatin. Previous studies have shown that these glutamic acid residues are crucial for Oct4’s ability to phase separate with MED1, both *in vitro* and *in vivo* ^[49]^, with mutations on those amino-acids limiting Oct4’s ability to induce cellular reprogramming^[52]^.

In summary, our data suggest that Oct4 clustering proceeds through a three-stage thermodynamic process. In the first stage, *nucleation*, Oct4–Oct4 self-interactions, driven primarily by non-ionic interactions among disordered regions, give rise to the formation of small clusters. In the second stage, *chromatin association*, multiple small Oct4 clusters anchor to chromatin through their DNA-binding domains, mediated largely by electrostatic interactions. By localising Oct4 clusters along the fibre, chromatin confines their free diffusion to a restricted spatial region. This confinement raises their local concentration and reduces the entropic cost of bringing multiple molecules together, thereby lowering the barrier for cluster growth. This step is governed primarily by the total amount of linker DNA available. In the third stage, *growth*, clusters expand through additional recruitment of Oct4 molecules to chromatin and by coalescence of chromatin-bound clusters. Here, linker length plays a central role: short linkers stabilise rigid fibres that restrict clusters to the chromatin surface, whereas long linkers confer higher flexibility, allowing Oct4 clusters to bend and loop the fibre, so that chromatin wraps around them, maximising DNA contacts and facilitating further growth. The coupling between enthalpic gain from multivalent Oct4–DNA and Oct4– Oct4 interactions, nucleosome–nucleosome contacts enabled by fibre looping, and the reduced entropic cost of coalescence, produces the largest Oct4 clusters at long linkers.

## Discussion

Our results reveal how the pioneer transcription factor Oct4 engages chromatin to remodel its structure and promote DNA accessibility. We find that Oct4 binds to chromatin and forms dynamic clusters that, in turn, transform chromatin structure. Importantly, Oct4 enhances DNA accessibility not by global decondensation, but by promoting nucleosome breathing and thereby driving chromatin into compact, irregular, and dynamic structures. These irregular structures are consistent with the liquid-like behaviour of chromatin triggered by nucleosome breathing^[26]^. In this liquid-like state, nucleosomes undergo frequent unwrapping and rewrapping at their entry–exit sites, which facilitates continuous nucleosome reorientation, neighbour exchange, and transient DNA exposure. Increasing Oct4 concentration amplifies this effect, while linker DNA length dictates how the liquid-like state emerges: medium and long linkers enable Oct4 to bridge distant nucleosomes and promote fibre condensation, whereas short linkers force Oct4 to intercalate between adjacent nucleosomes. Crucially, by amplifying DNA breathing, Oct4 suppresses the characteristic 10*n* versus 10*n* + 5 structural periodicity of chromatin fibres, replacing ordered zigzag geometries with heterogeneous inter-nucleosome interactions and driving all fibres towards a unified, liquid-like ensemble. Thus, Oct4 dynamically reshapes chromatin through a coupling between nucleosome breathing, cluster formation, and fibre mechanics, creating a structurally disordered yet accessible chromatin state.

In our simulations, we observe that formation of Oct4 clusters on chromatin proceeds through a three-stage thermodynamic process(Fig. 5):. In the first stage, nucleation, Oct4 molecules self-associate via their disordered regions to form small clusters. In the second stage, chromatin association, chromatin anchors Oct4 clusters via their DNA-binding domains. By localising Oct4 clusters to the fibre, chromatin confines their diffusion to a narrow region around the DNA, raising their local concentration and lowering the entropic penalty of bringing multiple molecules together. This step is governed primarily by the total amount of linker DNA available. In the third stage, growth, clusters expand through additional recruitment of Oct4 clusters and coalescence. Here, the linker length range is crucial due to its effect on chromatin mechanics: short linkers stabilise rigid fibres that restrict clusters to the fibre surface, whereas long linkers confer flexibility, allowing chromatin to bend and loop around the growing clusters, thereby maximising DNA contacts and promoting larger assemblies. This mechanism naturally couples Oct4 clustering with chromatin remodelling, bridging distal regions of the fibre and establishing a feedback loop between Oct4 organisation and chromatin structure.

**Fig. 5.**
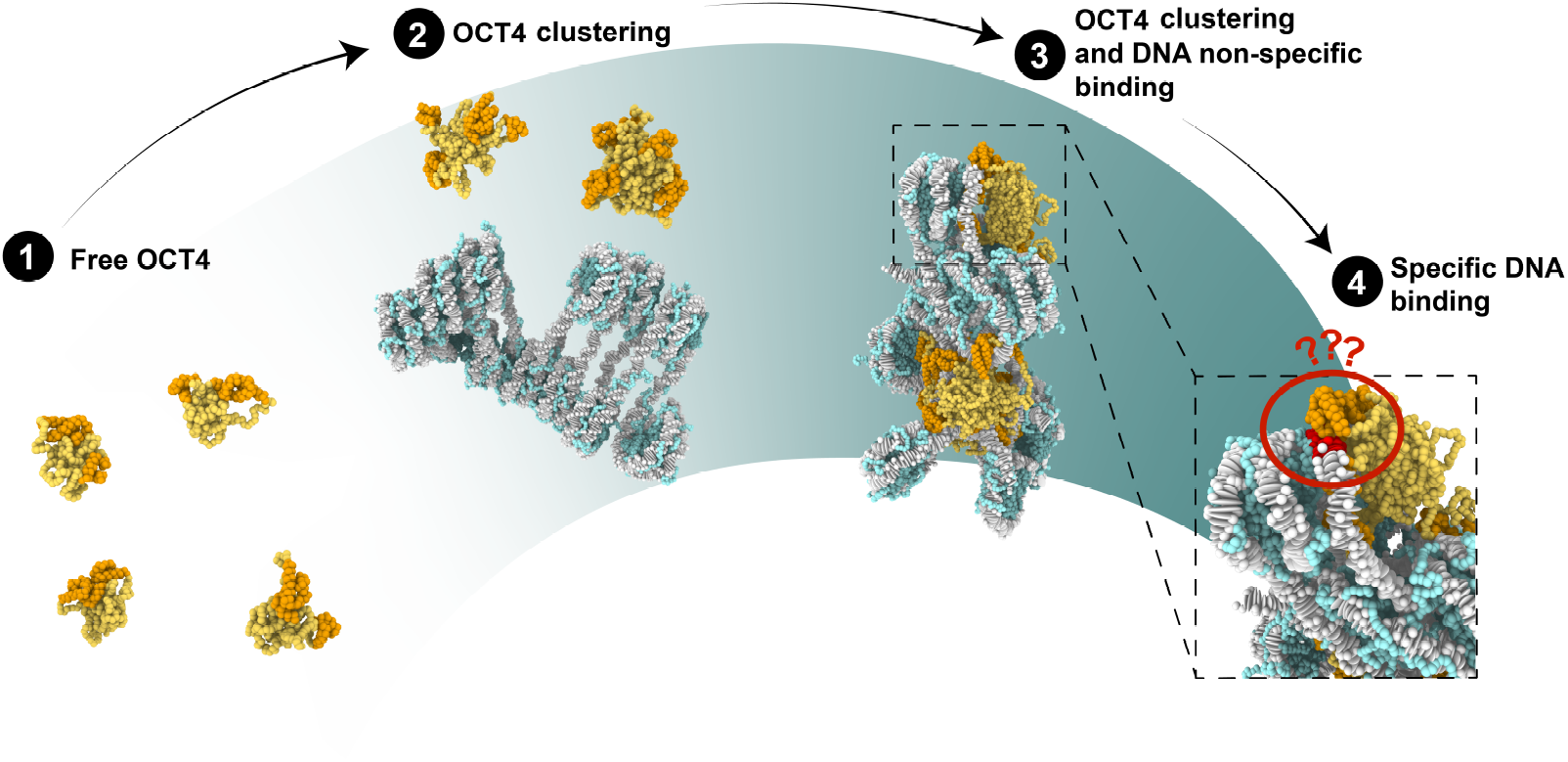
Proposed model for Oct4-mediated rearrangement of chromatin. We propose a model in which Oct4 forms clusters that become larger in the presence of chromatin. The binding of Oct4 clusters to chromatin reorganises chromatin structure, providing a mechanism by which Oct4 may target specific binding sites within closed chromatin.

This effect is reminiscent of threshold behaviours observed in complex coacervation, where heterotypic and homotypic multivalent interactions drive cooperative assembly. In this case, chromatin provides a flexible scaffold that facilitates clustering even at moderate Oct4 concentrations. Notably, the length of linker DNA controls the amount of exposed DNA, and thus the availability of anchoring sites for Oct4. While fine helical periodicity plays little role—because Oct4 promotes nucleosome breathing and overrides the inter-nucleosome orientations imposed by specific linker lengths—the overall linker length range determines whether chromatin behaves as a rigid or flexible fibre, thereby tuning Oct4’s capacity to grow large clusters. In agreement with prior evidence that linker DNA length modulates chromatin phase behaviour in vitro^[15,30,31]^, our results suggest that linker length may act as an additional layer of regulation, biasing transcription-factor binding and clustering towards specific genomic contexts. It should be noted that the Oct4:nucleosome stoichiometries in our simulations are higher than those typically observed physiologically. However, in reprogramming contexts where Oct4 is overexpressed, local concentrations may approach such ranges within particular genomic regions.

We further hypothesise that Oct4 clustering on chromatin, and the associated deformations that Oct4 clustering induces in chromatin, may help circumvent the so-called ‘search problem’: the paradox of how transcription factors locate their specific sites within the vast nuclear genome. During the early stages, chromatin anchors multiple Oct4 clusters in close proximity, effectively increasing their local concentration and facilitating encounters among them. As clusters grow, chromatin fibres bend and loop around Oct4 clusters, spatially confining distal genomic regions and allowing Oct4’s DNA-binding domains to explore a restricted neighbourhood more efficiently. Although our current model does not include base-specific interactions, such a mechanism could provide a biophysical explanation for how Oct4 accelerates the discovery and activation of silenced pluripotency genes during reprogramming. Future work incorporating explicit sequence-specific binding and epigenetic modulation of accessibility will be important to test this hypothesis.

Finally, our proposed mechanism likely acts in concert with other regulatory factors. Cooperative binding with Sox2, for example, is required for activation of Oct–Sox target genes, and enhances reprogramming efficiency^[71]^, and has been shown to play a role in single nucleosome recognition^[68,72]^, and in the recruitment of Oct4 to multi-protein condensates^[73]^. Different transcription factors may exploit distinct cluster-formation strategies: some may not form clusters at all, while others may require the presence of transcriptional co-activators acting as hubs for the recruitment of transcriptional machinery. Conversely, chromatin is not merely a passive scaffold: chromatin structure itself regulates the stability and material properties of phase–separated condensates^[15,31]^.

Overall, our work presents a biophysical mechanism in which Oct4 forms clusters on chromatin through a three-stage process of nucleation, chromatin association, and growth. This coupling of multivalent protein–DNA anchoring with activation-domain-mediated Oct4–Oct4 interactions facilitates chromatin reorganisation, potentially contributing to the resolution of the transcription factor search problem. These findings advance our mechanistic understanding of how transcription factors exploit multivalency and chromatin flexibility to sculpt chromatin architecture and influence DNA target accessibility. The modulation of such clustering behaviour—through concentration, cofactors, or epigenetic modifications—may constitute an additional regulatory layer in dynamic processes such as cellular reprogramming and cell-fate transitions.

## Materials and Methods

### Chemically-Specific Chromatin Model

Chromatin was simulated using our chemically-specific coarse-grained model^[26]^. In this model, we use one bead per amino acid, with beads maintaining the amino acid-specific hydrophobicity, charge and size of their atomistic counterpart. The secondary structure of the histone globular domains is maintained using a Gaussian elastic network model, whereas the histone tails are treated as fully flexible polymers. The DNA is represented with a modification of the Rigid Base Pair model, with every base pair treated as an ellipsoid decorated with two phosphate charges. The mechanical sequence-dependent behaviour of the DNA is captured by parametrising the six intra-base pair displacements in helical space from large-scale atomistic simulations^[74]^.

The full-length Oct4 protein was consistently coarse-grained at a one-bead per amino acid resolution (Fig. 1B). The coarse-grained model of the full Oct4 protein was built based on the crystal structure of Oct4 bound as a homodimer to the PORE motif^[43]^ (PDB accession code: 3L1P). The intrinsically disordered domains were added using Modeller^[75]^: 100 models were generated using the “slow” optimisation protocol, followed by a “slow” MD-based refinement protocol, and the model with the lowest discrete optimised energy was selected. Coarse-graining was performed by centring each bead per amino acid on their C_*α*_ positions. The two halves of the DNA-binding domain, the POU_S_ and the POU_HD_ domains, were treated as independent rigid bodies using the “fix rigid” command from LAMMPS. The intrinsically disordered regions (the linker connecting the POU domains and the N- and C-terminal activation domains) were treated as fully flexible polymers.

### Molecular dynamics simulations

MD simulations were performed in LAMMPS^[76]^ (version 3 March 2020), using our custom code^[26]^. We investigated the impact of Oct4 on chromatin fibres comprising 12 nucleosomes with Nucleosome Repeat Lengths (NRL = 147 bp of nucleosomal DNA plus the specified linker DNA length) of 167, 172, 177, 182, and 200 bp. For each NRL, simulations were carried out with 12, 24, or 48 Oct4 molecules interacting with chromatin fibres in which nucleosomes were modelled in a breathing configuration—where the nucleosomal DNA is allowed to stochastically associate and dissociate from the histone core—to account for the ability of Oct4 to induce breathing, as reported previously^[38]^. As a control, chromatin fibres were also simulated in the absence of Oct4, with nucleosomal DNA constrained in a non-breathing configuration^[26]^; here, the inner 147 bp of nucleosomal DNA were permanently bound to the histone core by an elastic network model.

Simulations of the control chromatin fibres were performed using Debye-Hückel replica exchange, with 16 replicas for each simulation condition^[26]^. The simulations were run at a constant temperature of 300 K using the Langevin thermostat. The Debye-Hückel screening parameter *κ* was varied across the replicas, from 8 Å^-1^ to 15 Å^-1^, with spacings optimised to ensure an exchange rate near 0.3. All systems were simulated for a total of 3 *µ*s, with a timestep of 10 fs. The initial 500 ns were discarded from the analysis to allow for system equilibration.

All Oct4-containing systems were simulated using an unbiased MD simulation protocol, since changing the Debye-Hückel screening parameter would result in the unbinding of Oct4 from the DNA, an interaction driven by electrostatics. For each system, 10 independent trajectories of 1 *µ*s were simulated to ensure convergence and representative sampling. The first 200 ns of each replica were discarded to allow the system to equilibrate.

All the visualisations and renderings were done using OVITO^[77]^.

### Analyses

#### Sedimentation coefficient

The sedimentation coefficient was computed using a custom Python implementation of the HullRad method^[78]^.

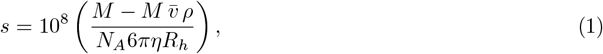

where *M* is the total molar mass, 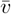 is the partial specific volume, *ρ* is the solvent density, *η* is the solvent viscosity, and *R*_*h*_ is the hydrodynamic radius. The coordinates of only the DNA beads were used to compute the convex hull of the structure,

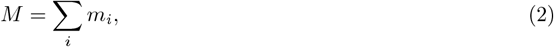

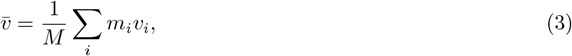

where *m*_*i*_ and *v*_*i*_ are the residue masses and specific volumes. The hydrodynamic radius *R*_*h*_ was calculated from the convex hull of the chromatin beads, expanded by a 2.8 Å hydration shell. Output values are reported in Svedberg units (S).

#### K-th nucleosome interactions

For the analysis of nucleosome neighbour interactions, as well as interaction orientation, the centre of mass (CoM) of the histone core, and its superhelical axis were computed for each nucleosome. Inter-CoM distances below 110 Å were counted as a nucleosome-nucleosome in interaction counts.

For every nucleosome pair, both the inter-CoM vector (*r*) and the super-helical axis (*z*) were used to compute their relative orientations. For every pair of nucleosomes *i* and *j*, we computed three angles defined as:

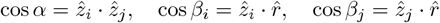

Pairwise interactions were then classified based on the following algorithm:

**if** *α <* 45^*°*^ **or** *α >* 135^*°*^:

**if** *β*_*i*_ *<* 45^*°*^ **or** *β*_*i*_ *>* 135^*°*^ **or** *β*_*j*_ *<* 45^*°*^ **or** *β*_*j*_ *>* 135^*°*^:

Face-face

**else:**

Side-side

**else:**

Face-side

#### Contact analysis

A contact was defined as any pairwise interaction where an atom of a given residue was found within 7.5Å of any atom belonging to a different molecule and falling within a specified partner category. To efficiently identify spatial neighbours, we used the “*cKDTree*” class from the “*scipy.spatial* “ Python module, which provides fast nearest-neighbour searches. Contacts were then normalised by frame and number of molecules.

To account for nucleosomal breathing and sliding, each DNA base was assigned to either nucleosomal or linker regions based on its local environment: bases that were in contact with histone core particles or located within ten base pairs of a contacted base were classified as nucleosomal, whereas those flanked by uncontacted stretches were defined as linker. This spatial classification allowed each Oct4 molecule to be categorised according to which DNA regions it contacted—nucleosomal, linker, or entry-exit (contacting both).

#### Cluster analysis

Cluster size analysis was performed using the OVITO Python API. Molecules were assigned to the same cluster if they were within 5.5 Å of each other. The resulting cluster sizes were extracted and normalised by the number of atoms in Oct4.

## Acknowledgements

We thank Sy Redding and Bryan Gibson for insightful discussions. JH is supported by the Herchel Smith Postdoctoral Fellowship Fund, and the EPSRC [grant number EP/X02332X/1] under the UKRI Post-doctoral Fellowships Guarantee Scheme [project TF-CHROM-LLPS]. This work was funded by a grant from the European Research Council (ERC) under the European Union’s Horizon 2020 research and innovation programme; 803326 to RCG. RCG and MJM are also supported by the UK Research and Innovation (UKRI) Engineering and Physical Sciences Research Council (EPSRC) under the UK Government’s guarantee scheme (grant EP/Z002028/1 awarded to RCG). MJM would like to acknowledge the Winton Programme for Physics of Sustainability for doctoral funding. This project utilised time on HPC resources granted by the UK High-End Computing Consortium for Biomolecular Simulation, HECBioSim (http://hecbiosim.ac.uk), supported by EPSRC (grant no. EP/X035603/1) to RCG.

## Notes

### Competing Interest Statement

The authors have declared no competing interest.

