## Supplementary Information for "Oct4 clusters promote DNA accessibility by enhancing chromatin plasticity"

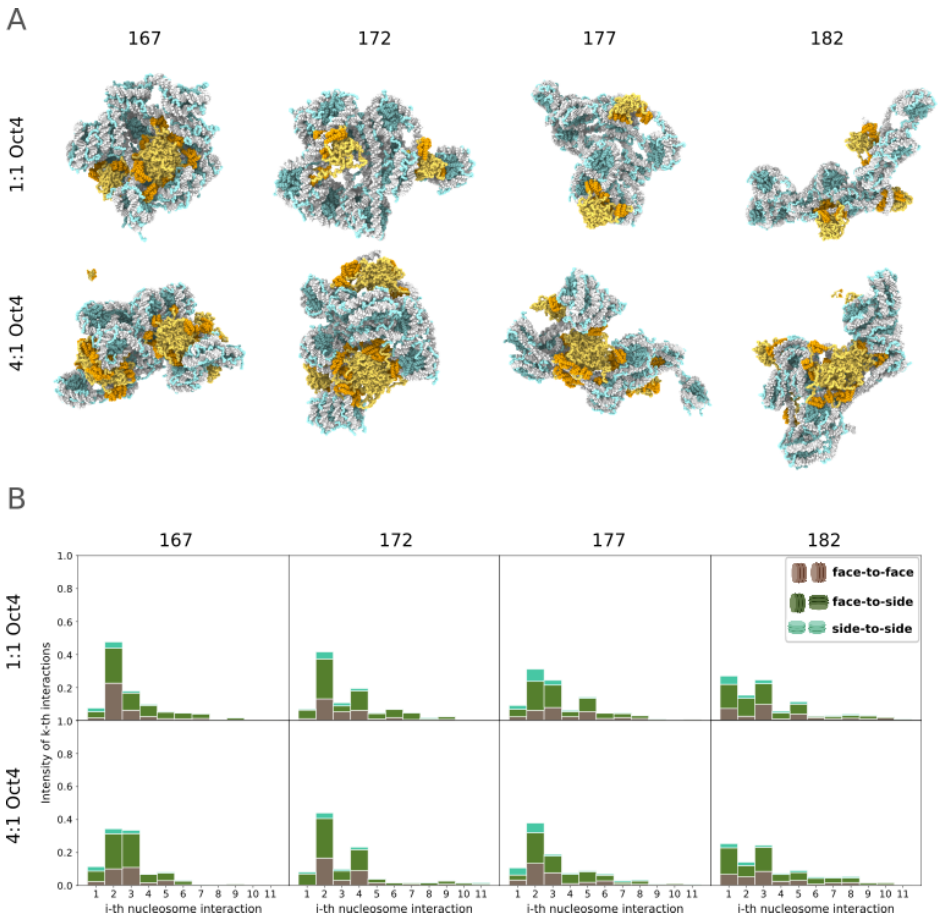


**Figure S1: Oct4-dependent chromatin compaction in different Oct4:nucleosome stoichiometries A.** Representative snapshots of chromatin with varying NRL, in the 1:1 (top row) and 4:1 (bottom row) Oct4:nucleosome stoichiometry. **B.** Frequency of interactions among k-th nearest nucleosome neighbours for chromatin with different NRLs, in the 1:1 and 4:1 Oct4:nucleosome stoichiometry. The bars are colour-coded to represent the percentage of nucleosome pairs involved in different types of inter-actions: face-to-face (brown), face-to-side (dark green), and side-to-side (turquoise). *See also Figure 2.*

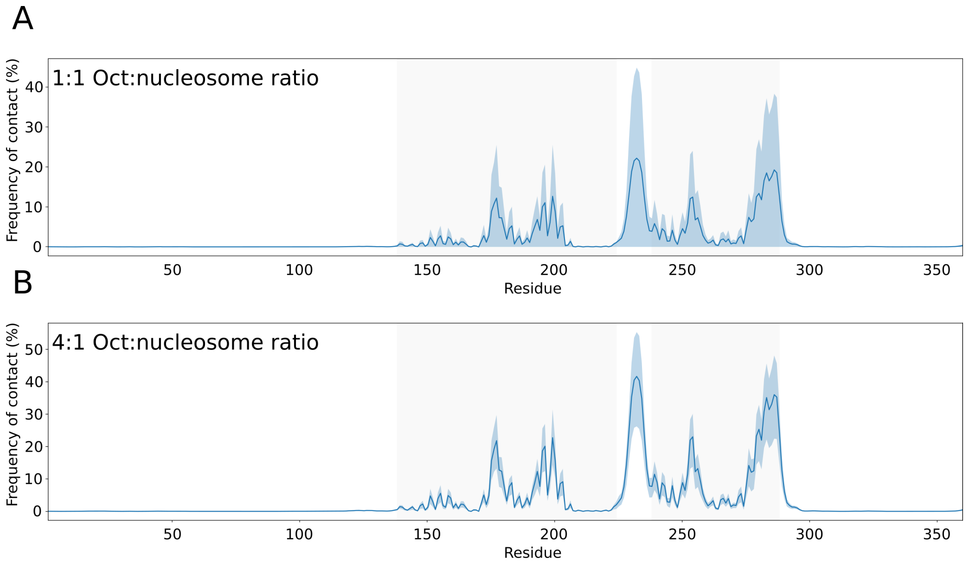


**Figure S2: Contact map of the Oct4-DNA interactions. A.** Per residue contact map of the Oct4-DNA interactions, averaged across all NRLs in simulations with 1:1 Oct4 nucleosome stoichiometry. **B.** Same as A, for simulations with 4:1 Oct4 nucleosome stoichiometry. *See also Figure 3.*

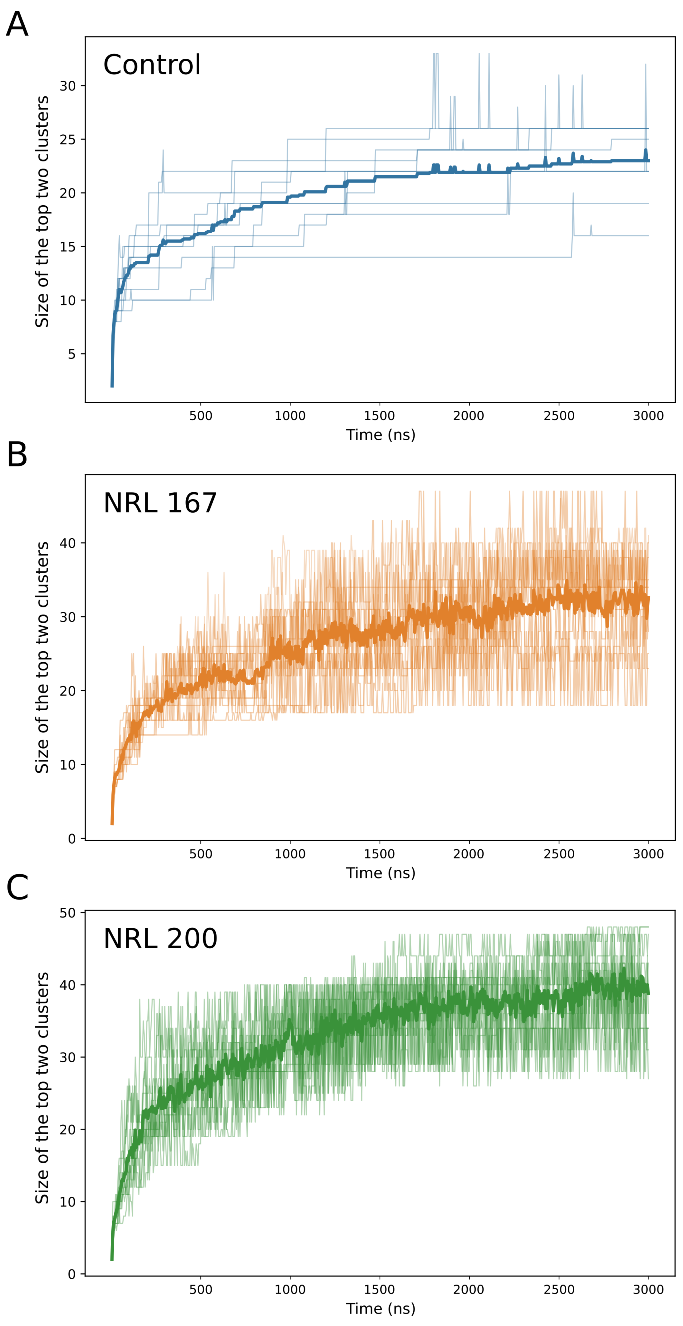


**Figure S3: Evolution of the size of Oct4 clusters. A.** Evolution of the sum of the size, in number of molecules, of the top two clusters, across 3 microseconds, for simulations without chromatin. In solid, the average of 10 trajectories is painted, whereas the individual trajectories are displayed at a lower opacity. **B.** Same as A, for simulations with chromatin with NRL 167. **C.** Same as A, for simulations with chromatin with NRL 200 *See also Figure 4.*

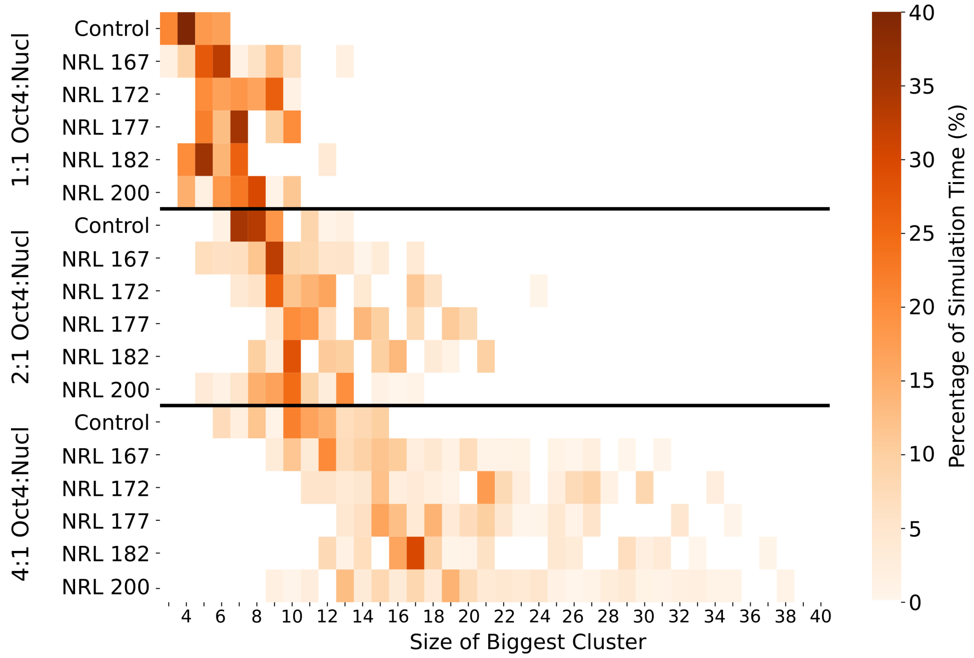


**Figure S4: Oct4 clusters are bigger in the presence of chromatin** Distribution of the size of the biggest cluster across the simulation. For each simulation condition, the percentage of time in which the biggest cluster has a given size is shown. *See also Figure 4.*

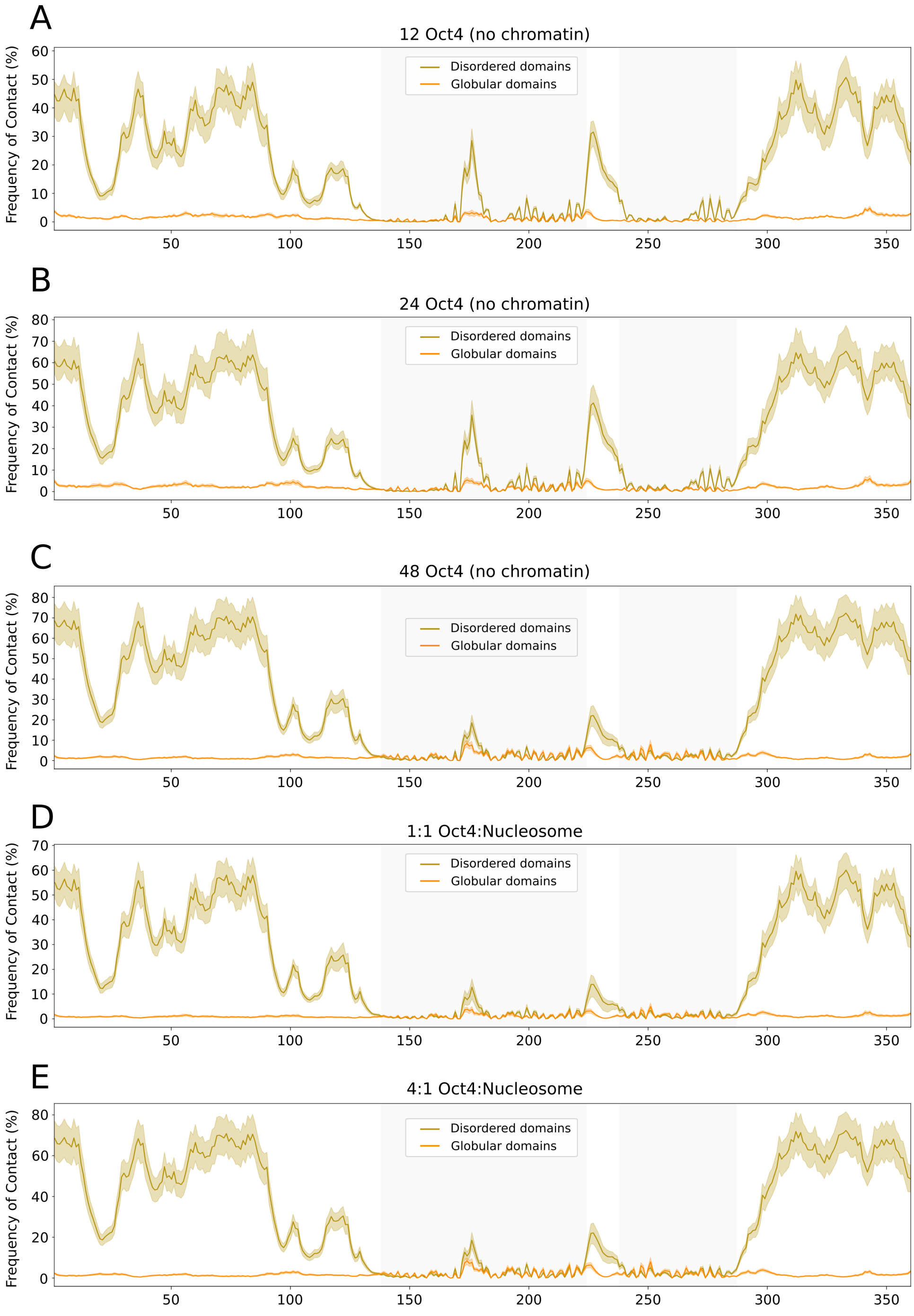


**Figure S5: Oct4 forms clusters via IDR contacts A.** Per residue contact map of the Oct4-Oct4 interactions, in simulations with 12 Oct4 copies in the absence of chromatin. **B.** Same as A, for simulations with 24 Oct4 copies in the absence of chromatin. **C.** Same as A, for simulations with 48 Oct4 copies in the absence of chromatin. **D.** Per residue contact map of the Oct4-Oct4 interactions, averaged across all NRLs in simulations of 1:1 Oct4 nucleosome stoichiometry. For each residue, the percentage of interaction with a residue belonging to a disordered domain or to a globular domain is depicted. The shadowed area represents the standard deviation. Contacts with disordered domains are represented in gold, and with the globular domains, in orange. **E.** Same as A, for simulations with chromatin with NRL 200. *See also Figure 4.*

**Supplementary Document 1**. Sequences used in the study

> **Oct4** (Human) – Uniprot *Q01860*

MAGHLASDFAFSPPPGGGGDGPGGPEPGWVDPRTWLSFQGPPGGPGIGPGVGPGSEVWGIPP
CPPPYEFCGGMAYCGPQVGVGLVPQGGLETSQPEGEAGVGVESNSDGASPEPCTVTPGAVKL
EKEKLEQNPEESQDIKALQKELEQFAKLLKQKRITLGYTQADVGLTLGVLFGKVFSQTTICR
FEALQLSFKNMCKLRPLLQKWVEEADNNENLQEICKAETLVQARKRKRTSIENRVRGNLENL
FLQCPKPTLQQISHIAQQLGLEKDVVRVWFCNRRQKGKRSSSDYAQREDFEAAGSPFSGGPV
SFPLAPGPHFGTPGYGSPHFTALYSSVPFPEGEAFPPVSVTTLGSPMHSN

**>NRL_167**

ACTTACATGCACAGGATGTAACCTGCAGATACTACCAAAAGTGTATTTGGAAACTGCTCCAT
CAAAAGGCATGTTCAGCTGGATTCCAGCTGAACATGCCTTTTGATGGAGCAGTTTCCAAATA
CACTTTTGGTAGTATCTGCAGGTGATTCTCCAGGGCGGCCAGT

**>NRL_172**

AGTACTTACATGCACAGGATGTAACCTGCAGATACTACCAAAAGTGTATTTGGAAACTGCTC
CATCAAAAGGCATGTTCAGCTGGATTCCAGCTGAACATGCCTTTTGATGGAGCAGTTTCCAA
ATACACTTTTGGTAGTATCTGCAGGTGATTCTCCAGGGCGGCCAGTAC

**>NRL_177**

TAAGTACTTACATGCACAGGATGTAACCTGCAGATACTACCAAAAGTGTATTTGGAAACTGC
TCCATCAAAAGGCATGTTCAGCTGGATTCCAGCTGAACATGCCTTTTGATGGAGCAGTTTCC
AAATACACTTTTGGTAGTATCTGCAGGTGATTCTCCAGGGCGGCCAGTACTTA

**>NRL_182**

ATGTAAGTACTTACATGCACAGGATGTAACCTGCAGATACTACCAAAAGTGTATTTGGAAAC
TGCTCCATCAAAAGGCATGTTCAGCTGGATTCCAGCTGAACATGCCTTTTGATGGAGCAGTT
TCCAAATACACTTTTGGTAGTATCTGCAGGTGATTCTCCAGGGCGGCCAGTACTTACA

**>NRL_200**

TCCTGTGCATGTAAGTACTTACATGCACAGGATGTAACCTGCAGATACTACCAAAAGTGTAT
TTGGAAACTGCTCCATCAAAAGGCATGTTCAGCTGGATTCCAGCTGAACATGCCTTTTGATG
GAGCAGTTTCCAAATACACTTTTGGTAGTATCTGCAGGTGATTCTCCAGGGCGGCCAGTACT
TACATGCACAGGAT
